# Growth Dynamics of Ductal Carcinoma in Situ Recapitulate Normal Breast Development

**DOI:** 10.1101/2023.10.01.560370

**Authors:** Marc D. Ryser, Matthew A. Greenwald, Inmaculada C. Sorribes, Lorraine M. King, Allison Hall, Joseph Geradts, Donald L. Weaver, Diego Mallo, Shannon Holloway, Daniel Monyak, Graham Gumbert, Shariar Vaez-Ghaemi, Ethan Wu, Kevin Murgas, Lars J. Grimm, Carlo C. Maley, Jeffrey R. Marks, Darryl Shibata, E. Shelley Hwang

## Abstract

Ductal carcinoma in situ (DCIS) and invasive breast cancer share many morphologic, proteomic, and genomic alterations. Yet in contrast to invasive cancer, many DCIS tumors do not progress and may remain indolent over decades. To better understand the heterogenous nature of this disease, we reconstructed the growth dynamics of 18 DCIS tumors based on the geo-spatial distribution of their somatic mutations. The somatic mutation topographies revealed that DCIS is multiclonal and consists of spatially discontinuous subclonal lesions. Here we show that this pattern of spread is consistent with a new ‘Comet’ model of DCIS tumorigenesis, whereby multiple subclones arise early and nucleate the buds of the growing tumor. The discontinuous, multiclonal growth of the Comet model is analogous to the branching morphogenesis of normal breast development that governs the rapid expansion of the mammary epithelium during puberty. The branching morphogenesis-like dynamics of the proposed Comet model diverges from the canonical model of clonal evolution, and better explains observed genomic spatial data. Importantly, the Comet model allows for the clinically relevant scenario of extensive DCIS spread, without being subjected to the selective pressures of subclone competition that promote the emergence of increasingly invasive phenotypes. As such, the normal cell movement inferred during DCIS growth provides a new explanation for the limited risk of progression in DCIS and adds biologic rationale for ongoing clinical efforts to reduce DCIS overtreatment.

## INTRODUCTION

Mammography screening has been successful in reducing breast cancer mortality,^1-3^ yet its benefits are accompanied by harms such as false positive findings, unnecessary procedures, and overdiagnosis.^4^ The overdiagnosis of indolent tumors that would not cause any harm in the woman’s remaining lifetime is of particular concern for patients diagnosed with ductal carcinoma in situ (DCIS).^5^ DCIS is considered a precursor of invasive breast cancer, yet studies support that as many as 70-80% of DCIS found on mammography would not progress to invasive cancer if left untreated.^6,7^ Because it is currently not possible to accurately distinguish indolent from aggressive DCIS, nearly all DCIS patients undergo surgery, and many receive additional radiation and endocrine therapy.^8^ This strategy leads to widespread overtreatment, affecting as many as 40,000 women each year in the US alone.^9^

The goal of breast cancer screening is to intercept the progression from normal breast tissue to invasive cancer. Current dogma purports that this transformation occurs in a linear stepwise fashion, with DCIS being a proximate step before invasion.^7^ Indeed, DCIS and invasive breast cancer share similar morphologic, proteomic and genomic alterations,^10-13^ and frequently the only histologic distinction between DCIS and invasive cancer is abnormal tumor cell migration beyond the basement membrane. Given the genomic similarity between DCIS and invasive breast cancer and the ability of DCIS to spread within the ductal tree over several centimeters, one might expect that abnormal cell movement is an inherent feature of DCIS growth. However, there is a lack of evidence to support this claim, and the common observation of “skip” lesions with large segments of intervening normal tissue within DCIS is not explained by the current model.

In animal models, cell lineage markers can be traced in situ to reconstruct epithelial breast cell movement.^14^ In such studies, individual progenitor cells are labeled in vivo and the migration of their progeny (subclones) is inferred from the final topographic distribution of lineage markers. This approach has been used to quantify the dynamics of murine pubertal breast duct development, whereby ducts grow, branch, and penetrate the surrounding stroma through a process called branching morphogenesis (**Figure 1**).^15-17^ Importantly, normal duct growth does not occur by continuous subclone spreading but is orchestrated by advancing growth buds that each contain multiple stem cell subclones.^15^ These stem cells intermittently contribute to ductal growth, leading to multiclonal ducts whose subclones form spatially discontinuous ‘skip’ patterns.

**Figure 1:**
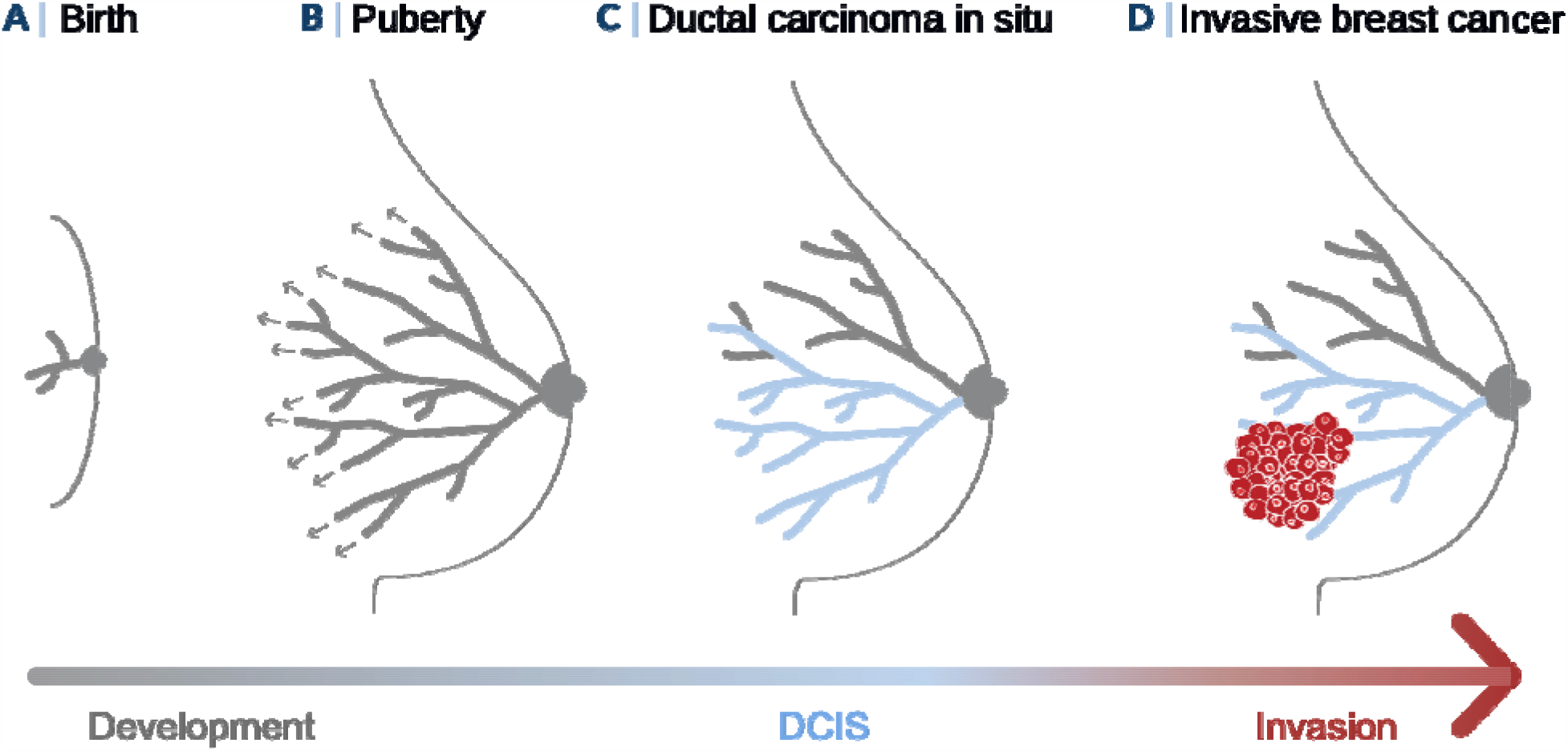
The female breast: From normal development to invasive cancer. **(A)** At birth, the mammary gland consists of the simple embryonic rudiment. **(B)** During pubertal development, the embryonic rudiment undergoes branching morphogenesis and develops into the adult ductal tree. **(C)** Ductal carcinoma in situ (DCIS) consists of neoplastic cells that are contained within the ducts and lobules of the adult mammary gland. **(D)** During invasive progression, DCIS cells penetrate the basement membrane of the ducts and lobules and invade the breast stroma.

While direct observation of cell movement in human breast epithelium is impractical, somatic mutations uniquely label the progeny of individual subclones.^18^ We thus postulate that cell migration in human DCIS can be inferred from the spatial distribution of somatic mutations. We reconstruct the three-dimensional mutation topographies of 18 DCIS tumors over macroscopic length scales of up to 7cm and find many spatially discontinuous subclones that are difficult to reconcile with canonical clonal evolution.^19^

Interestingly, a model of DCIS growth that mimics the dynamics of branching morphogenesis of the normal breast (**Figure 1**) naturally recapitulates the discontinuous mutation patterns observed in DCIS. We propose that normal cell movement conferred by branching morphogenesis-like growth reveals a biological basis for why many DCIS grow to a macroscopic size and then remain stable for decades without progression to invasive cancer.

## RESULTS

### Multiregional sequencing reveals spatial mutation topographies

We identified 18 women who had undergone surgery for a diagnosis of screen-detected DCIS, including 9 patients with DCIS tumors alone (*pure* DCIS), and 9 patients with DCIS tumors adjacent to invasive breast cancer (*synchronous* DCIS). All tumors were of nuclear grade 2 or 3 and most (14/18) were hormone-receptor positive. The most common histologic patterns were solid and cribriform type, and most tumors (15/18) exhibited comedo-like features **(Suppl. Table S1)**. From each surgical specimen we obtained between 2 and 5 spatially separated formalin-fixed and paraffin-embedded (FFPE) tissue regions, and in each tissue section we microdissected^20^ and spatially registered small regions, or spots, each containing approximately 100 to 500 epithelial cells **(Figure 2A, Suppl. Figure S1)**. In addition to spots containing individual ducts with DCIS, we microdissected normal breast ducts, ducts with benign breast disease, and areas of synchronous invasive cancer. To complement the spatial and histologic spot annotations, we determined the genotype of each spot through targeted sequencing of tumor-specific mutation panels derived from whole exome sequencing (WES) of macro-dissected DCIS foci.

**Figure 2:**
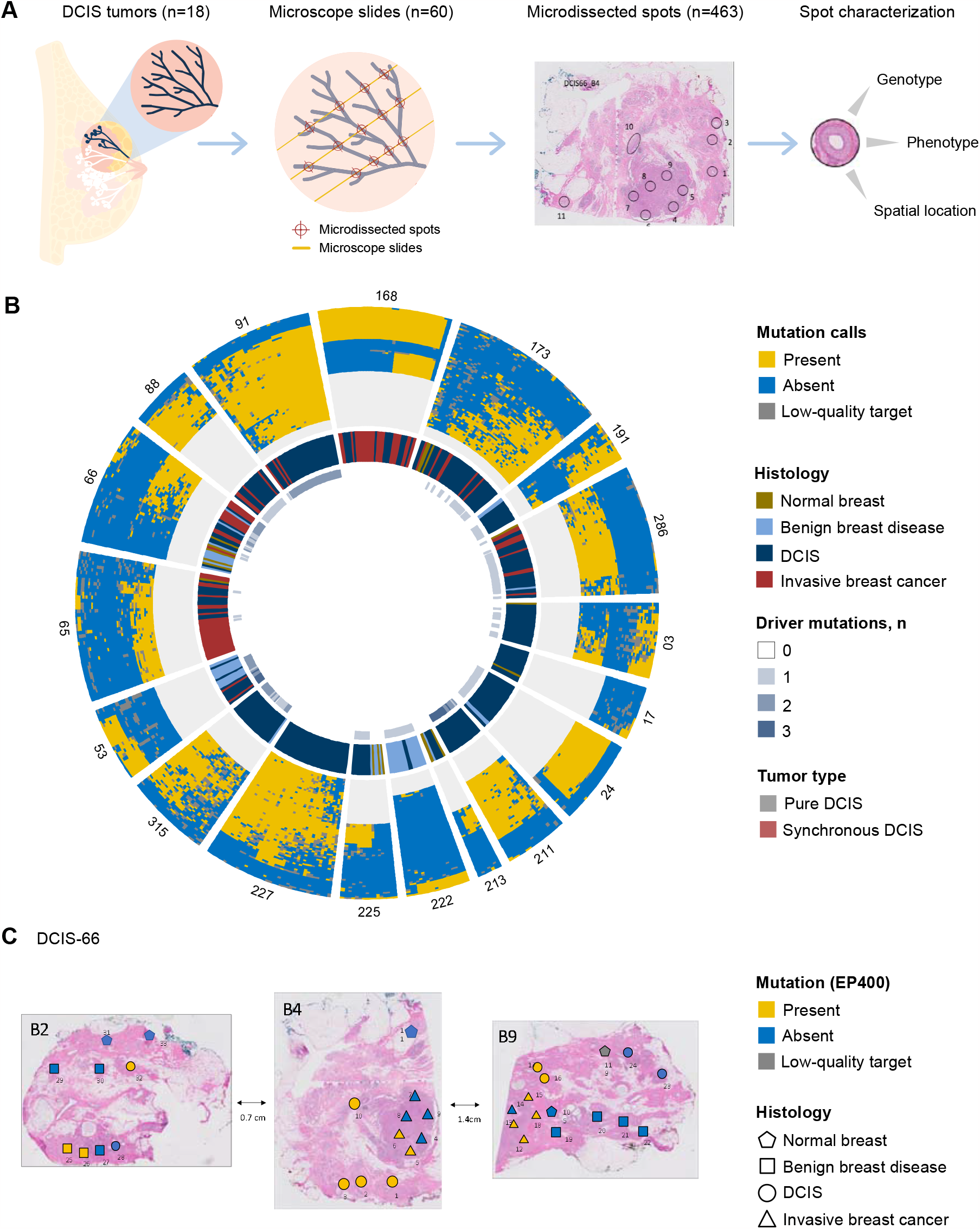
Multiregional sequencing reveals spatial mutation topographies of DCIS tumors. **(A)** Between 2 and 5 spatially separated microscope sections were obtained from 18 DCIS tumors. From each microscope slide, small tissue areas (spots) were microdissected, spatially registered, histologically annotated (normal breast duct, duct with benign breast disease, duct with DCIS, invasive breast cancer), and genotyped. Genotyping was based on targeted sequencing of tumor-specific mutation panels that had been derived from whole exome sequencing analyses of macro-dissected DCIS areas. **(B)** Summary of the genetic and phenotypic spot data for all 18 DCIS tumors. Each sector groups together spots of the same tumor, and tumor labels are shown at the periphery. Differences in height of the outermost track (mutation calls) reflect the varying mutation panel sizes for each tumor. **(C)** Spatial pattern of a select mutation in DCIS-66 (gene: EP400, chr12:132472310). Shapes indicate spot histology and colors the mutation status.

After eliminating germline mutations and low-quality targets **(Suppl. Figure S2**), the final study cohort comprised 463 individual spots across 60 tissue sections (**Suppl. Table S1**). The resulting dataset (**Figure 2B**) combined phenotypic and genotypic annotations of the spatially registered spots. In addition to 313 spots with DCIS, we registered 87 spots with invasive cancer, 46 spots with benign breast disease, and 17 spots with normal breast ducts, all confirmed by pathology review. A total of 823 (median per tumor: 45, range: 24-66) mutation targets were identified by WES, of which 558 (68%; median per tumor: 31, range: 8-59) mutations were detected by targeted sequencing (**Suppl. Figure S3**). Across all 558 mutations we identified two de novo mutational signatures that matched established consensus signatures implicated in carcinogenesis (**Suppl. Figure S4**). Across the 18 DCIS tumors we identified a total of 21 putative driver mutations (median per tumor: 1, range: 0-3) (**Suppl. Table S2**). Combining the genotypic spot characterizations with the spatial tumor maps, we constructed geospatially annotated somatic mutation topographies for each DCIS (**Figure 2C**).

### DCIS is a multiclonal and heterogeneous disease

The resulting spatial-genetic data were used to characterize the clonality and intratumor heterogeneity (ITH) of the DCIS portions within each tumor. Indeed, the variant allele frequencies (VAFs) of somatic mutations within individual spots contain valuable information about the structure of cell populations in the local cellular neighborhood (**Figure 3A**). Variant allele frequencies of 50% or greater reflect locally clonal mutations that are present in all cells of the sampled duct cross-section, whereas VAFs below 50% indicate locally subclonal mutations carried by a subpopulation of resident cells only. Across the 313 histologically confirmed DCIS spots in our cohort, the within-spot VAF spectra of detected mutations were generally subclonal and dispersed, as evidenced by low median values and high inter-quartile ranges, respectively (**Figure 3B**). These data demonstrate that most DCIS ducts contain an admixture of distinct genetic subclones which vary in frequency throughout the lesion. This finding of multiclonal DCIS is consistent with previous single cell-based studies.^11,19,21^

**Figure 3:**
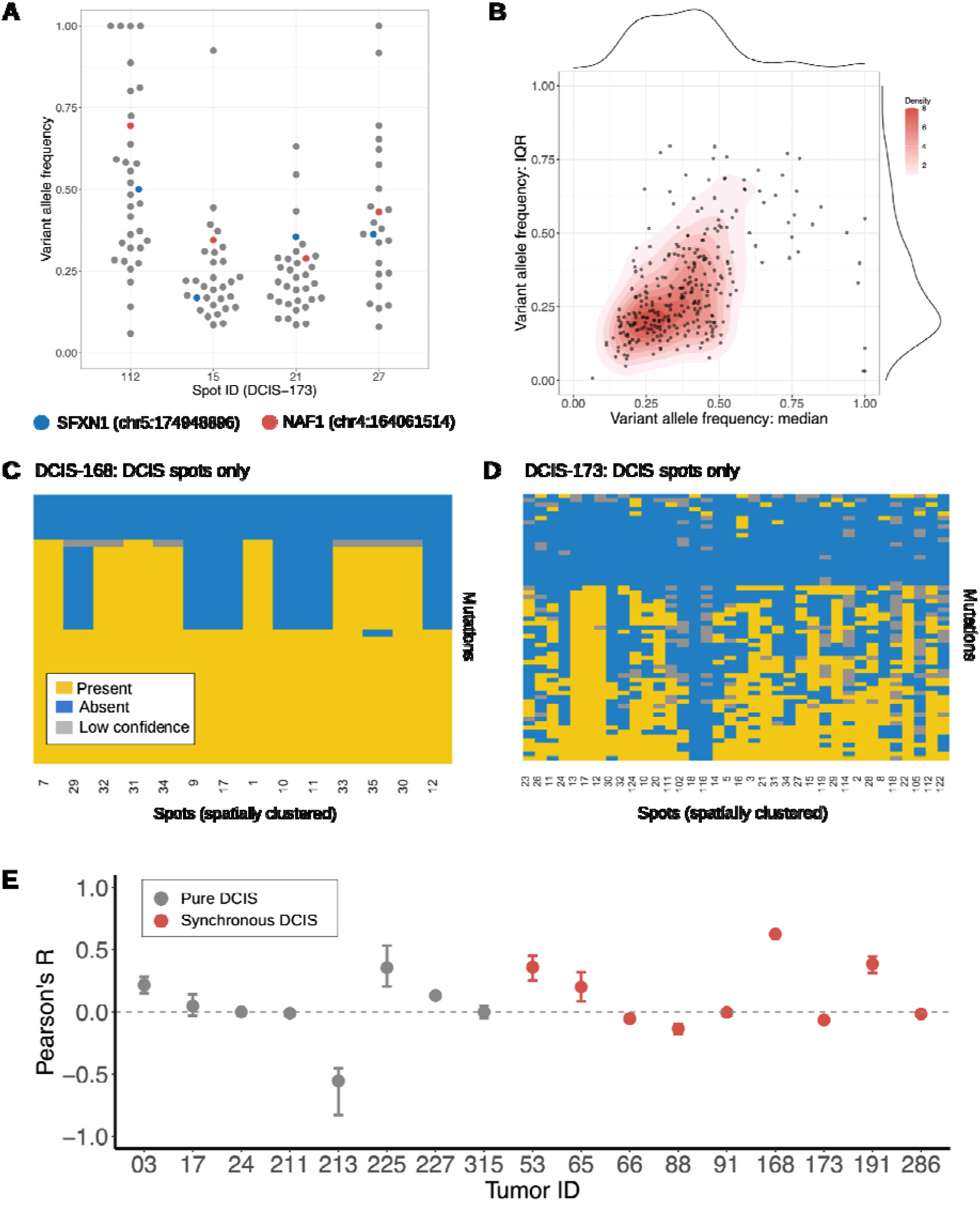
DCIS tumors are multiclonal and spatially heterogeneous. All analyses in this figure are restricted to DCIS spots. **(A)** The variant allele frequency (VAF) spectra of detected mutations are shown for 4 select spots in DCIS-173; the VAF of two select mutations in the genes SFXN1 (blue) and NAF1 (red) are highlighted. **(B)** Bivariate summary statistics for spot-level VAF spectra are shown across all DCIS spots (n=313) of the 18 tumors, with median V F on the x-axis, and interquartile range (IQR) of the VAF on the y-axis. Red color scheme visualizes spot density. **(C)** Mutation patterns for all DCIS spots in DCIS-168 are organized by hierarchical clustering of mutations (rows) and spatial clustering of spots (columns); spatial clustering was based on one-dimensional t-distributed stochastic neighbor embedding (t-SNE) of the spots’ spatial coordinates. **(D)** Mutation patterns for all DCIS spots in DCIS-173, see panel C for details and color legend. **(E)** For each tumor, the spatial correlations of DCIS spot genotypes were quantified using Pearson’s R; DCIS-222 was excluded because it had only 2 DCIS spots. Monte Carlo sampling was used to account for posterior uncertainty of mutation calls, resulting in predicted means (circles) and 95% prediction intervals (bars). Median predicted mean correlation was -0.01, without detectable differences between pure DCIS and synchronous DCIS with adjacent invasive cancer (p=.81, Wilcoxon rank-sum test).

To quantify the degree of genetic ITH, we defined spot genotypes as the binary vectors of somatic mutation calls (present/absent), visualized as the columns of the mutation panels (**Figure 3C-D, Suppl. Figure S5**). While some DCIS tumors comprised only few distinct spot genotypes (e.g., **Figure 3C**), most contained a substantial number of distinct genotypes (e.g., **Figure 3D**), which is indicative of pervasive ITH. Notably, we observed a lack of spatial clustering of similar spot genotypes (**Figure 3C-D, Suppl. Figure S5**), suggesting limited spatial correlations of duct genotypes. We further investigated this by computing the correlations of spatial and genetic spot distances (**Figure 3E**) and found that most tumors exhibited low spatial-genetic correlations (median: -.01), without detectable differences between pure and synchronous DCIS tumors (p=.81, Wilcoxon rank-sum test).

In summary, these findings support the presence of multiclonal ducts and extensive spatial heterogeneity within each DCIS tumor,^11-13,19^ but do not address when and how such ITH arises during tumorigenesis. To investigate this, we turned our attention to the spatial topographies of individual somatic mutations.

### Expansive skip lesions favor a model of early evolution

We categorized mutations as *public* (present in ≥90% of DCIS spots in the tumor) or *restricted* (present in <90% of DCIS spots); the latter are particularly informative because they allow for tracking of individual subclones in space. Across the 17 tumors with more than 2 DCIS spots, we identified a total of 379 restricted mutations (**Suppl. Table S1**). Interestingly, restricted mutations often spanned expansive but discontinuous tumor regions of up to 7cm in diameter, and in 14 of 17 tumors, one or more restricted mutations covered the entire DCIS portion (**Figure 4A**). This finding of expansive mutational skip lesions is consistent with two recent studies that performed spatial subclone mapping in DCIS tumors.^19,22^

**Figure 4:**
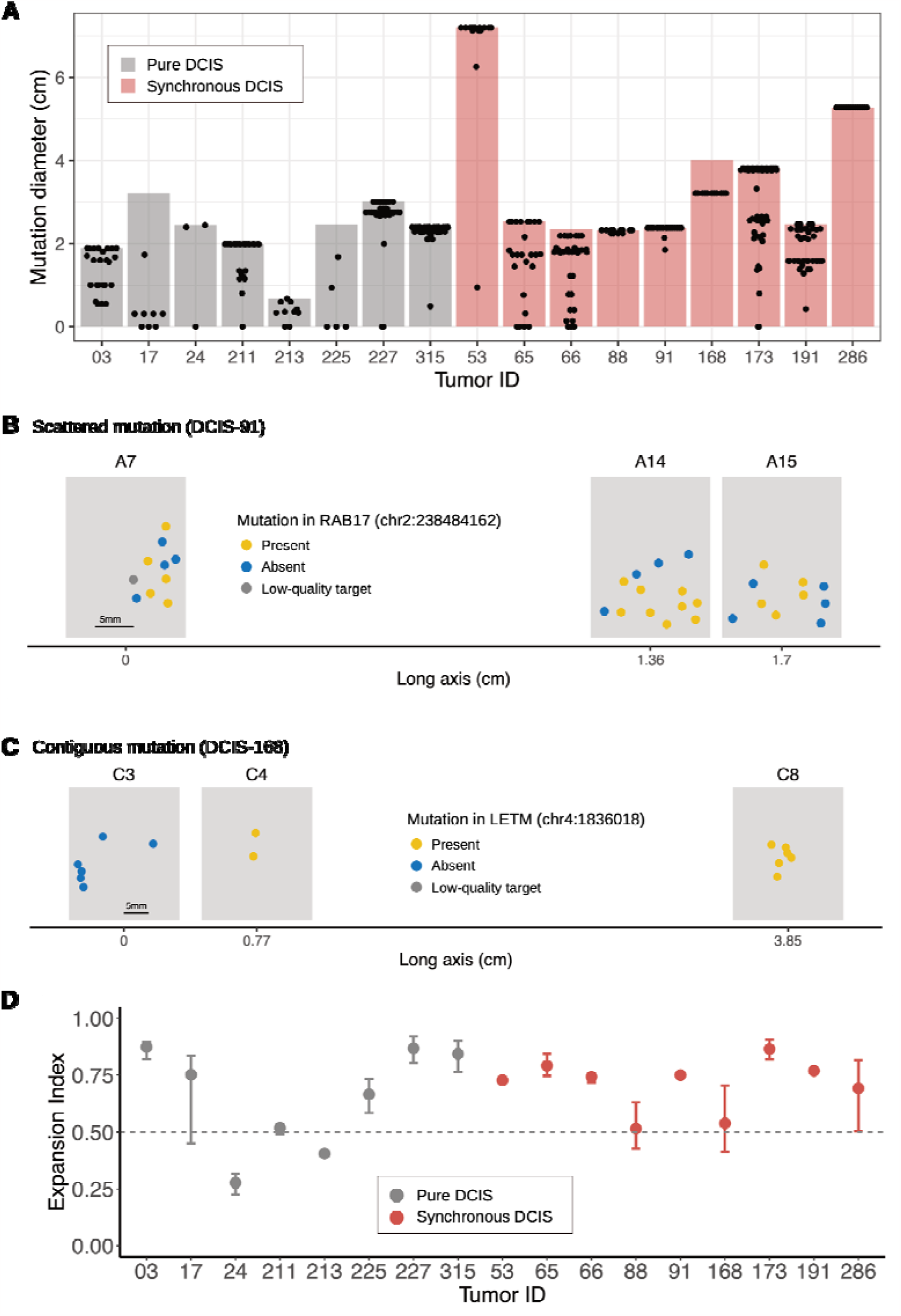
DCIS mutations form expansive skip lesions. All analyses in this figure are restricted to DCIS spots; DCIS-222 was excluded because it only had 2 DCIS spots. **(A)** The diameter of restricted mutations (found in <90% of spots; black dots) relative to the extent of the DCIS tumor itself (bar). **(B)** Scattered mutations are characterized by a lack of spatial separation between spots that do and do not contain the mutation. An example from DCIS-91 is shown. Grey rectangles represent the microscope sections (x-y plane) along the tumor’s long (z-) axis. **(C)** Contiguous mutations are characterized by a spatial separation of spots that do and do not contain the mutation. An example from DCIS-168 is shown; see also description of panel B. **(D)** The expansion index (EI) of a tumor characterizes the degree of mutation scattering, ranging from contiguous ( ) to scattered ( ). Monte Carlo sampling was used to account for posterior uncertainty of mutation calls, resulting in predicted means (circles) and 95% prediction intervals (bars). Median EI was 0.74 across all tumors, without detectable difference between pure DCIS (median: 0.71) and synchronous DCIS with adjacent invasive cancer (median: 0.74; p=0.88, Wilcoxon rank-sum test).

Mutational skip lesions can arise by two distinct mechanisms, depending on whether evolution takes place early or late in the growth process. In the early evolution scenario, subclonal mutations arise during the early expansion from the first DCIS cell and then disperse across the ductal tree during expanding tumor growth. In the late evolution scenario, the mutations arise late during tumor expansion and disseminate across the tree through extensive sweeps, in competition against less fit subclones.

Delineation of these two scenarios is possible because they predict different types of spatial mutation patterns. In the early evolution scenario, the passive dissemination of early mutations is expected to produce scattered mutation topographies, or ‘skip’ lesions (**Figure 4B**). In contrast, in the late evolution scenario, late mutations that expand through subclonal sweeps are expected to produce more contiguous mutation patches (**Figure 4C**). To test these predictions against the data, we introduced a new tumor-level measure, the expansion index (EI), which ranges from 0 to 1 and measures whether a lesion is dominated by disperse (*EI* ≫ .5) or contiguous (*EI* ∼ .5) mutations (**Methods** and **Suppl. Figure S6**). The median EI across all tumors was 0.74, and 12/17 (71%) tumors had an EI in the disperse range of, *EI* ≥ .6 (**Figure 4D**). Notably, there was no detectable difference in EI between pure DCIS (median: 0.71) and synchronous DCIS (median: 0.74, Wilcoxon rank-sum test: p=0.88). The consistently elevated expansion indices are indicative of mutational skip lesions and suggest that the widespread ITH is likely due to the passive dissemination of early subclones in the early evolution scenario.

Two additional observations provide evidence against the late evolution scenario of mutation dissemination. First, expansive subclonal sweeps are expected to yield locally homogeneous ducts,^23^ which is at odds with the observation of subclonal VAFs at the spot level (**Figure 3B, Suppl. Figure S7**).^19^ Second, expansive subclonal sweeps would require the acquisition of a substantial cellular fitness, yet we only found a limited number (n=21) of putative driver mutations in our cohort (**Suppl. Table S2**), and there was no evidence that driver mutations were more disperse than passenger mutations (**Suppl. Figure S8**).

In theory, copy number changes producing spatially localized losses of mutant alleles can account for discontinuous mutation patterns. In practice, however, such a mechanism would need to be very pervasive to account for the widespread skip lesions in our data. To ascertain the likelihood that copy number aberrations formed the primary mechanism for discontinuous mutation patterns, we performed spatial copy-number profiling across 19 spots of a large DCIS tumor in our cohort. Copy-number profiles across DCIS ducts were stable (**Suppl. Figure S9**), and in spots where both copy number and DNA mutation data were available, none of the absent mutations coincided with an allelic loss (**Suppl. Figure S9**).

In summary, our data support a model of early evolution where genetic subclones arise during the initial expansion from the first DCIS cell and before dispersion across the ductal tree through expansive tumor growth. What remains unclear, however, are the cellular mechanisms that govern this expansive growth phase.

### DCIS growth recapitulates normal ductal morphogenesis

The scattered mutation topographies we inferred from our data (**Figure 5A**) are strikingly analogous to patterns observed during the normal pubertal development of murine mammary ductal trees.^15,17^ During breast development, individual mammary stem cells contribute to ductal expansion only intermittently to produce dispersed subclone patterns along the branching ductal tree. Based on these similarities, we posited the ‘Comet model’ of DCIS tumorigenesis which recapitulates the stochastic fate rules of ductal elongation and binary branching as inferred from pubertal branching morphogenesis.

**Figure 5:**
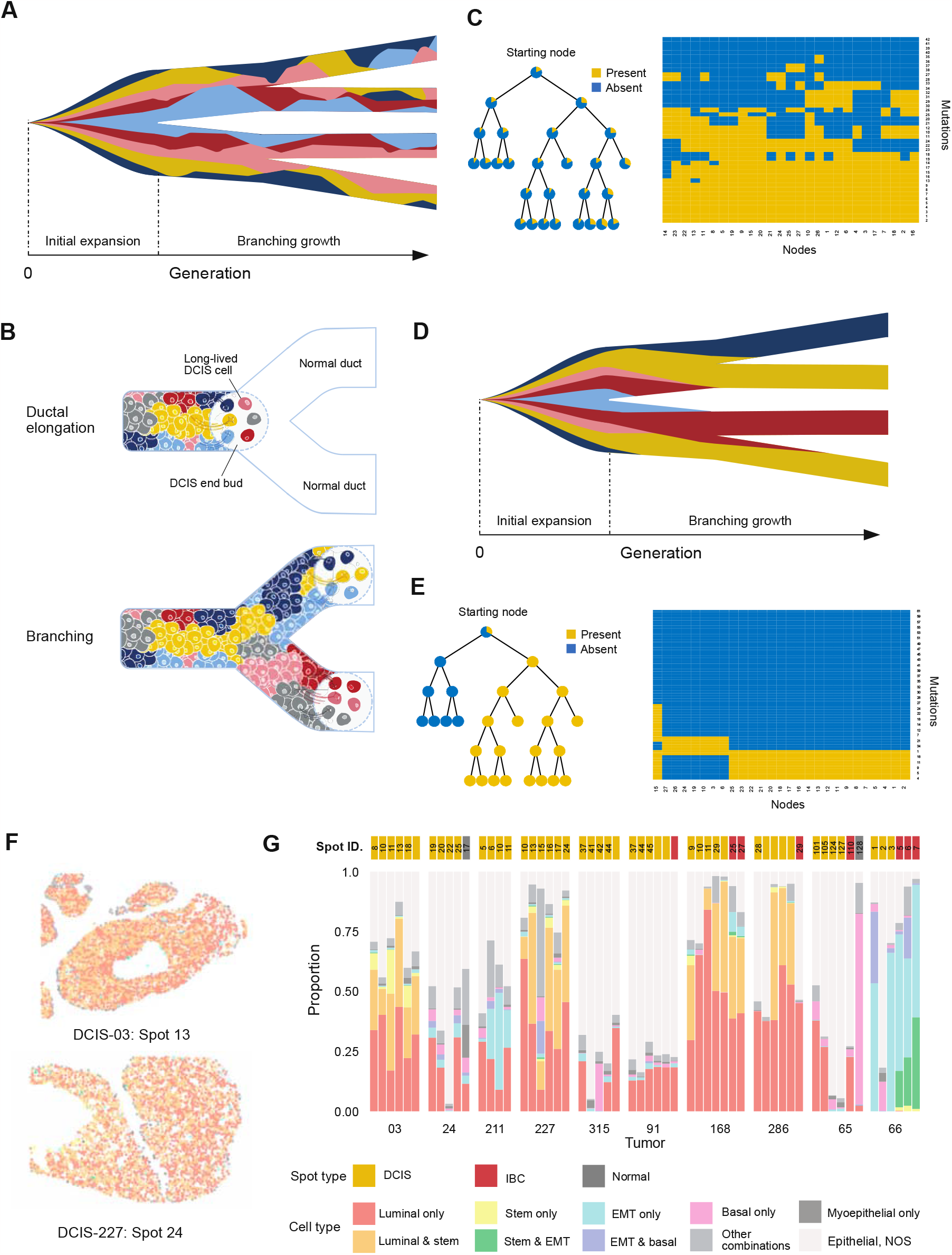
The Comet model of DCIS tumorigenesis. **(A)** A modified Muller plot illustrating the typically observed data in our cohort. After initial expansion of early subclones, the growth patterns are characterized by multiclonal ducts and disperse skip lesions. **(B)** The Comet model of DCIS growth recapitulates the dynamics of pubertal branching morphogenesis. During ductal elongation (top), the long-lived neoplastic cells of the DCIS end bud undergo intermittent proliferation; after transit-amplification, the clustered progenies of the long-lived cells become embedded in the growing multiclonal DCIS duct. During branching (bottom), the end bud cells are randomly distributed between the two daughter branches where they duplicate, and the two resulting end buds start growing along their respective daughter branches. **(C)** Mutation patterns resulting from the Comet model. Left: DCIS growth is initiated at the starting node and propagated across the ductal tree, with pie charts indicating the local variant allele frequencies (VAFs) of a select mutation. Right: the hierarchically clustered mutation pattern corresponding to the simulation in the left panel, illustrating the local presence/absence of mutations (rows) across the examined duct cross-sections (columns). **(D)** A modified Muller plot illustrating the expected subclone frequencies that arise from a canonical model of cancer evolution along the ductal tree. Initial expansion of the first DCIS cell and subsequent branching growth are governed by quasi-neutral clonal evolution. Due to the thin tube-like geometry of the ducts, individual subclones are expected to rapidly go extinct or fixate, resulting in monoclonal ducts. **(E)** As in C, but instead using a canonical model of cancer evolution, see Methods for details. **(F)** Spatial distribution of epithelial cell types in two DCIS-filled ducts, generated by multiplexed ion-beam imaging (MIBI). Each field of view (FOV) is of size 500 x 500 ; corresponding color legend at the bottom of panel G. **(G)** A total of 57 FOVs across 10 tumors, including 49 DCIS ducts, 2 normal breast ducts, and 8 areas of invasive cancer were analyzed using MIBI. Where applicable, spot ID (top) maps each FOV to the corresponding spot label from the mutational analysis. Epithelial cells (PanCK+) were classified as either luminal (BCL2+ and/or GATA3+), stem-like (PAX5+ and/or SOX10+), basal (CK5+), epithelial-to-mesenchymal (Vimentin+), or myoepithelial (SMA+); for cells assigned to multiple subtypes, we distinguished the three most common combinations, and grouped the less frequent combinations; cells that did not match any of the subtypes were classified as not otherwise specified (NOS).

The Comet model posits that DCIS growth is driven by the expanding end buds of the tumor front which contain populations of long-lived neoplastic cells that arise early in evolution (**Figure 5B, Methods**). These long-lived cells stochastically undergo episodic expansion to produce the subclone populations that populate the elongating DCIS duct. When an expanding tumor bud reaches a branching point in the ductal tree, the long-lived cells are randomly divided between the two daughter ducts and then duplicate. Such comet tail-like backward seeding of subclones naturally results in multiclonal DCIS ducts and expansive mutational skip lesions across the involved portions of the mammary tree. Simulations of the Comet model illustrate the expansive dispersion of subclonal mutations and high levels of ITH (**Figure 5C**).

On the other hand, because DCIS shares many morphologic, proteomic, and genomic features with invasive breast cancer,^10-13^ it would appear natural for its growth to be governed by the uncontrolled cellular proliferation and subclone competition of canonical clonal evolution. Yet when combined with the branching topology and thin tube-like geometry of the ductal tree, these dynamics are expected to result in rapid stochastic fixation or extinction of individual mutations along the ductal tree^23^ (**Figure 5D**). Indeed, simulations indicate a smaller number of subclones and limited ITH (**Figure 5E**) when compared to the Comet model (**Figure 5C**).

To quantify the ability of the Comet model to explain the spatial-genetic data in our cohort, we developed a computational platform that mimics our experimental design (**Methods** and **Technical Appendix**). In brief, we generated a stochastic ductal tree in silico, randomly seeded the first tumor cell, simulated the DCIS growth dynamics, and recorded the simulated VAFs of sampled DCIS ducts in the final tumor. We fit the model to the experimental data using approximate Bayesian computation (ABC) and found that it agreed with salient summary statistics of the empirical mutation topographies (**Suppl. Figure S10A**). Through formal Bayesian model selection, we showed that the Comet model provided a superior fit compared to a model of clonal evolution, as evidenced by a Bayes’ factor^24^ of 11.7 (**Suppl. Table S3, Suppl. Figure S10-B**).

In summary, these data support a novel Comet model of DCIS growth, whereby genetic heterogeneity is acquired early and multiple subclones are disseminated across the ductal tree through a process that recapitulates the branching morphogenesis of normal pubertal breast development. This model not only provides a simple explanation for the observed discontinuous mutation patterns in DCIS,^19^ but also generates testable hypotheses regarding the local composition of DCIS ducts.

### Stable hierarchical cell populations

The Comet model posits that local DCIS cell populations are deposited during growth by stochastically expanding progenitor cells. As with normal gland development, these clonal subpopulations are expected to be maintained by a stable hierarchical mixture of progenitor, transit-amplifying, and more mature luminal cells.^15,16^ To test this hypothesis we characterized the local epithelial subtype compositions of 57 individual spots from 10 tumors in our cohort through multiplexed ion-beam imaging (MIBI).^25^ Using a machine learning algorithm, individual epithelial cells were classified as stem-like, basal, luminal, epithelial-to-mesenchymal (EMT), or myoepithelial (**Figure 5F**), thus allowing us to characterize the local cell type composition in each spot (**Figure 5H**). As predicted by the Comet model, individual DCIS ducts consistently comprised a hierarchical mixture of more differentiated luminal cells and less differentiated stem-like and basal cells.

We further performed targeted DNA methylation sequencing of individual DCIS ducts from 6 tumors in our cohort (**Suppl. Figure S11**). We found extensive epigenetic diversity, which further corroborates the notion that DCIS ducts are maintained by a stable epithelial hierarchy rather than clonal competition and frequent subclonal sweeps.

### Phenotypic plasticity and multiclonal invasion

Phenotypic heterogeneity is common in DCIS^13,26,27^ and may be driven by the underlying genotypic heterogeneity. However, because it has been difficult to map mutations to phenotypes,^13,27^ phenotypic heterogeneity may also be the result of phenotypic plasticity, whereby cells of the same genotype express different phenotypes in response to their local microenvironment. In our cohort, we found evidence of phenotypic plasticity in the form of many shared mutations between spots with benign breast disease, DCIS and invasive cancer (**Figure 6A-B, Figure 6D-E, Suppl. Figure S12**). To further investigate potential plasticity, we focused on the 8 synchronous DCIS tumors with adjacent invasive cancer.

**Figure 6:**
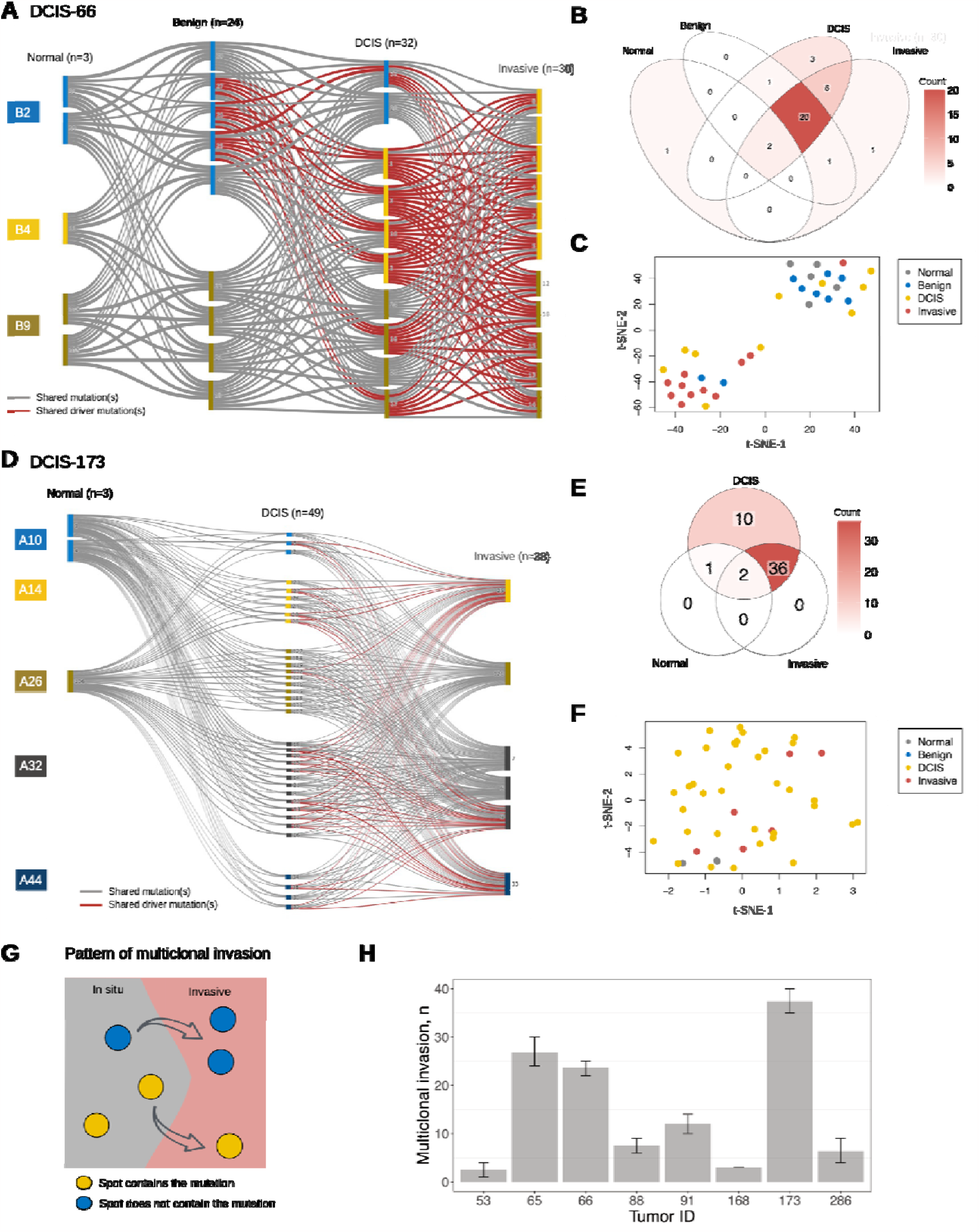
Phenotypic plasticity and multiclonal expansion. (A-C) Mutational summary of DCIS-66. (A) Mutation flow across the phenotypic spectrum of breast disease, from normal breast tissue and benign breast disease to DCIS and invasive cancer; n indicates the total number of mutations detected among spots of a given histology. The vertical rectangles represent individual spots, and their color indicates the corresponding microscope slide. Grey connections indicate one or more shared mutation(s) in the absence of shared putative driver mutations, and red connections indicate one or more shared putative driver mutation(s). **(B)** Venn diagram summarizing shared mutations (drivers and passengers) across spot histologies. **(C)** t-distributed stochastic neighbor embedding (t-SNE) of spot genotypes, with colors indicating spot histology. **(D-F)** Mutational summary of DCIS-173. See captions of panels A, B, and C for details about panels D, E and F, respectively. **(G)** Example of a mutation pattern that indicates multiclonal invasion: the mutation is present in some but not all DCIS spots, and in some but not all invasive spots. Such a pattern indicates that two distinct cell populations, one with and one without the mutation, are present both inside and outside the ducts. **(H)** Multiclonal invasion patterns were found in all 8 tumors that had both DCIS and invasive spots; duplicate patterns were excluded. Monte Carlo sampling was used to account for posterior uncertainty of mutation calls, resulting in predicted means (bars) and 95% prediction intervals (error bars).

Most somatic mutations were shared between in situ and invasive spots (mean: 89%, range: 78-100%), and among the 3 tumors that also contained ducts with benign breast disease, a substantial fraction of mutations was shared across all three phenotypes (**Suppl. Table S4**). Putative driver mutations found in the invasive tumor portions were consistently present in adjacent DCIS and benign breast disease ducts. In genotype space, DCIS and invasive spots tended to co-cluster (**Figure 6C, Figure 6F, Suppl. Figure S12A-H**), and we observed genotypic co-clustering of all three phenotypes in 2 of 3 tumors with benign breast disease ducts (**Suppl. Figure S12B-C**). A similar lack of correlations between genotype and phenotype was observed with respect to the spots’ local cell type composition (**Suppl. Figure S13**). Taken together, a pervasive lack of phenotype-genotype correlations in these tumors suggests phenotypic plasticity.

Phenotypic plasticity can also result in multiclonal invasion, that is the co-migration of multiple genetic subclones from the ducts into the stroma as they encounter a permissive microenvironment.^11^ At the single mutation level, multiclonal invasion manifests itself in the form of mutations that are present in some but not all DCIS spots, and in some but not all invasive spots (**Figure 6G**). Counting the number of unique such patterns in each tumor, we found evidence of multiclonal invasion in all 8 synchronous tumors (**Figure 6H**). In addition to being multiclonal, invasion was spatially dispersed, with admixed clusters of in situ and invasive spots across multiple spatially separated sections in 7 of the 8 tumors (**Suppl. Figure S1**). In summary, the spatially disperse patterns of multiclonal invasion are consistent with phenotypic plasticity in DCIS.^11^

## DISCUSSION

Based on the mutation topographies of 18 human DCIS tumors, we propose the Comet model of DCIS tumorigenesis. The Comet model posits that multiple genetic subclones arise shortly after the first DCIS cell, and then disperse across the ductal tree through a mechanism that recapitulates the branching morphogenesis of normal breast development.

Because of its histologic and genomic similarity with invasive breast cancer, DCIS is often considered “just one step” away from invasion. Yet this characterization is at odds with a growing recognition that most DCIS tumors remain latent for decades if left untreated.^6,28^ The Comet model offers a potential solution to this clinical incongruency. Indeed, the branching morphogenesis of normal breast development is a regulated expansion where mobile progenitor cells proliferate, differentiate, and branch to form new ductal elements but remain confined within the basement membrane. By recapitulating this developmental program of mobile expansion, many DCIS tumors can grow into macroscopic yet stable neoplasms without reliance on the uncontrolled proliferation and abnormal mobility of invasive cancer. Importantly, in contrast to neoplastic growth governed by clonal evolution, the Comet dynamics are not subject to incessant subclone competition that produces increasingly aggressive phenotypes. The proposed model thus provides a simple explanation for the common occurrence of indolent DCIS tumors and provides biologic rationale for an evolving clinical paradigm that seeks to de-escalate treatment in low-risk DCIS patients.^5,29^

The Comet model is consistent with previously reported multiclonality and intratumor heterogeneity of DCIS tumors,^10-13^ and expands this knowledge with a novel explanation for the co-occurrence of duct-level multiclonality and global subclone dispersal. While the origins of multiclonality *per se* can be explained^11^ by an early punctuated burst of genomic instability,^30-32^ the simultaneous occurrence of duct-level multiclonality and global subclone dispersal have been difficult to reconcile.^19^ Indeed, under a canonical model of cancer growth—characterized by uncontrolled proliferation and clonal evolution—the thin tube-like mammary ducts are expected to accelerate local sweeps, resulting in contiguous patches of monoclonal DCIS ducts.^23^ This currently accepted model is at odds the observed mutational patterns, but is readily resolved by the proposed Comet model, where the tumor’s expanding end buds contain multiple, episodically proliferating subclones that produce local multiclonality and global subclone dispersal.

A subgroup analysis of 8 patients with DCIS and adjacent invasive cancer supported a multiclonal invasion model^11^ in which multiple subclones co-migrate from the ducts into the stroma. Such multiclonal invasion, taking place at spatially distant foci and amidst a paucity of putative driver mutations, could arise through convergent stepwise progression where each physical focus represents an independent evolutionary bottleneck. Yet the Comet model provides a more parsimonious scenario in which invasion is facilitated by a conducive local microenvironment rather than being conferred by accumulated somatic mutations. This model is strikingly consistent with the previously described plasticity of both normal breast tissue^33^ and DCIS tumors^27^, and suggests that certain DCIS tumors are essentially *born to be bad* and ready to invade when and where permissive conditions are met. More fundamentally, it remains unclear what differentiates indolent from progressive DCIS tumors, although recent studies suggest primary roles for the tumor microenvironment such as early changes in the ductal myoepithelium^25^ or the immune microecology.^34^Similar studies performed in colorectal cancers (CRC)^30,35,36^ provide a direct comparison of cancer growth patterns between the two organs. In both sites, growth is driven by long-lived progenitor cells, situated in the growing end buds of DCIS^15^ and at the base of CRC glands,^37^ respectively. Furthermore, the branching of DCIS ducts is analogous to the fission of cancer glands during CRC growth.^37^ Yet while the transit-amplifying progenies of CRC stem cells exit the gland within a few days, their DCIS counterparts are embedded in the expanding duct and provide a genomic record of the end buds’ proliferative activity during growth. This difference can explain why CRC glands are generally monoclonal populations dominated by a single fixated subclone, whereas duct cross-sections contain multiple subclones. This comparison highlights the likely role of tissue architecture in shaping the mode of evolution.^23,38,39^

Our study has limitations. First, due to sequencing constraints in FFPE samples, spot selection was biased toward larger ducts. While this may have led to an underestimation of overall heterogeneity, our findings of local multiclonality and global subclone dispersion would be invariant under the inclusion of smaller ducts. Second, because patient-specific mutation panels were derived from microdissected DCIS areas, they did not contain mutations private to the invasive compartment of synchronous tumors. While this may have led us to overestimate the fraction of mutations shared between DCIS and adjacent invasive cancer, our findings are consistent with a body of literature documenting the genomic similarity between DCIS and adjacent invasive cancer.^10-13^ Third, we cannot exclude the possibility that long-range seeding of individual cells may be responsible for the observed skip lesions. However, given the lack of evidence for such cellular migration–across macroscopic distances and through often densely packed DCIS ducts–the Comet model provides a more parsimonious explanation. Fourth, since our cohort was composed of intermediate to high grade and mostly hormone receptor positive tumors with solid or cribriform growth patterns, the Comet dynamics may not be applicable to other pathologic subtypes, such as micropapillary DCIS and low-grade tumors. Finally, while it is commonly assumed that DCIS cells grow along the pre-existing mammary ductal tree, an alternative model of neoductogenesis^40^ proposes that DCIS may branch off the pre-existing tree to grow its own subtrees. However, as long as the subtrees resulting from neoductogenesis are topologically invariant, our mathematical models remain applicable.

An expansive and structured penetration of the breast stroma in the absence of invasion and metastasis is an inherent feature of normal pubertal breast development. In this study, we provide evidence that DCIS cell migration recapitulates this developmental process of normal branching morphogenesis, resulting in indolent tumors that are susceptible to mammographic overdiagnosis. Interestingly, the process of branching morphogenesis is not unique to the breast and is equally implicated in the development of the prostate, thyroid, and lung.^17,41,42^ The intriguing observation that cancer overdiagnosis is common in these organs as well^43-45^ raises the possibility that a recapitulation of developmental branching morphogenesis could be a contributing factor to the etiology of indolent tumors across cancer sites.

## METHODS

### Patient cohort and biological samples

The study was approved by the Institutional Review Board of the Duke University Medical Center (protocol Pro00054877), and a waiver of consent was obtained according to the protocol. We identified patients diagnosed with screen-detected breast cancer who underwent breast-conserving surgery or mastectomy at Duke University Medical Center between 1999 and 2016. During the selection process, formalin-fixed paraffin-embedded (FFPE) tissue blocks for cases with a complete spatial block map were obtained from the Duke Pathology archives. Each block was pathology reviewed (A.H.) for diagnosis according to the WHO classification of tumors.^46^ A total of 21 cases with tumor tissue present in two or more FFPE blocks were identified through this process, including 11 patients with *pure DCIS* tumors, and 10 patients with DCIS tumors with synchronous ipsilateral invasive breast cancer (*synchronous DCIS*). DCIS nuclear grade and estrogen- and progesterone-receptor status were abstracted from the patients’ medical records. As described below, a total of 3 patients were excluded prior to final data analyses, because of technical issues (n=2) or insufficient information content (n=1). The final analytic cohort thus comprised 18 patients, 9 with pure DCIS and 9 with synchronous DCIS (**Suppl. Table S1**). Finally, we collected matched normal samples for all patients, in the form of blood (n=4), uninvolved lymph nodes (n=4), or adjacent, morphologically normal breast tissue (n=9).

### Whole exome sequencing

To design tumor-specific mutation panels, whole exome sequencing (WES) was performed on bulk tissue samples as follows. For each patient, two or more spatially separated (≥8mm) FFPE blocks were identified, and areas containing DCIS (but no invasive cancer) were macro-dissected from between 10 and 25 hematoxylin-stained tissue sections (5 microns thick). The first and last sections were stained with hematoxylin-eosin (H&E) and reviewed by a study pathologist (A.H.) to confirm that tumor cellularity was at least 70%. DNA was extracted using the FFPE GeneRead DNA Kit according to manufacturer instructions. DNA quantity was determined using a QubitTM 1X dsDNA HS Assay Kits (ThermoFisher, cat. n. Q33230), and DNA quality was assessed using the Agilent 2100 Bioanalyzer. WES was performed on ≥40ng of genomic DNA from each sample. Each aliquot was sheared to a mean fragment length of 250 bp (Covaris LE200), and Illumina sequencing libraries were generated as dual-indexed, with unique bar-code identifiers, using the Accel-NGS 2S PCR-Free library kit (Swift Biosciences, cat. n. 20,096). We pooled groups of 96 equimolar libraries (100 ng/library) for hybrid capture of the human exome as well as a targeted panel of the exons of 83 breast cancer genes, using IDT’s xGen Exome Research Panel v1.0; see Fortunato et al.^47^ for details. After hybridization, capture pools were quantitated via qPCR (KAPA Biosystems kit), and the final product was sequenced using an Illumina HiSeq 2500 1T instrument (multiplexing nine tumor samples per lane). After binning the data based on its index identifier and aligning it to the Genome Reference Consortium Human Build 37 (GRCh37) using the BWA-MEM algorithm,^48^ sequencing duplicates were identified using Picard’s MarkDuplicates (GATK). The resulting BAM files were then used to design the tumor-specific mutation panels as described in the next section. The WES protocol was performed at the McDonnell Genome Institute at Washington University School of Medicine in St Louis.

### Tumor-specific mutation panels

For each patient, we designed a tumor-specific target panel of single nucleotide variants (SNVs) based on the BAM files obtained from WES of tumor and matched normal tissue. Variants were called using the software MuTect^49^ (Broad Institute), using default settings. Starting from a combined set of SNVs that had “judgment=KEEP” in at least one of the two samples, we excluded SNVs not mapped to chromosomes 1 through 22 or the X chromosome, SNVs identified as single nucleotide polymorphisms in dbSNP^50^ and SNVs that were within 300bp of another SNV. For patients where more than 100 SNVs remained after these exclusions, we decreased the final panel size to 100 or less by first removing mutations at a variant allele frequency (VAF) below 10% in both bulk samples and taking a simple random sample if necessary. SNVs identified in COSMIC^51^ (https://cancer.sanger.ac.uk/cosmic) were included independently of the above filter settings.

### Saturation microdissection

From each tumor, between 2 and 5 spatially separated FFPE blocks that contained individual DCIS ducts or lobules suitable for microdissection were identified by the study pathologists (AH, DS). In mixed tumors, the study pathologists (A.H. and D.S.) further identified DCIS-adjacent areas of IBC suitable for microdissection. From each block, between 5 and 10 consecutive 5-micron tissue sections were prepared on plastic slides and lightly stained with H&E. A study pathologist (DS) then microdissected small tissue areas, or spots, using selective ultraviolet light fractionation (SURF) as previously described^52^ and implemented by our group.^36^. In brief, a micromanipulator was used to place small ink dots over individual duct cross-sections and, in the case of synchronous DCIS tumors, over equivalently sized areas of invasive breast cancer. The absolute number of tumor cells in each microdissected spot was estimated to be between 100 and 500 cells. After the destruction of unprotected DNA through 3-4 hours of short-wave ultraviolet light irradiation, individual ink dots were removed from the slides using a pipette tip and placed in a microfuge tube for DNA extraction.

### Targeted mutation sequencing

After proteinase K and TE treatment at 60°C for 4 hours, and then at 98°C for 10 minutes, AMPure XP beads (Beckman Coulter) were added (1.2x) to extract the DNA. Polymerase chain reaction (PCR) was performed directly on the dried beads (35-40 cycles) using a custom AmpliSeq primer for the tumor-specific SNV panels as described above. PCR repeatedly failed for two tumors and led to their exclusion from further analysis (DCIS-118, DCIS-158). Barcoded libraries (One-step, Qiagen) were then run on MiSeq or NextSeq Illumina sequencers, with an average coverage of >500x and a minimum coverage of 20x for each mutation. The FASTQ files from the sequencers were uploaded to the Galaxy web platform and analyzed using the public server (http://usegalaxy.org/).^53^ Briefly, our Galaxy pipeline included FASTQ grooming, adapter trimming (TrimGalore), short read alignment (BWA) to GRCh37, Naive Variant Caller and Variant Annotator. For each locus, we defined the reference and alternate alleles based on the WES results and recorded their respective read counts from the targeted sequencing runs.

### Targeted methylation sequencing

Custom AmpliSeq primers were designed for a target amplicon (chromosome 7; positions 77395824 to 77295930) containing 5 consecutive CpG sites, as well as three amplicons with LUMP (leukocytes unmethylation for purity) sites to assess epithelial content (cg10559416, cg21376733, cg27215100).^54^ After proteinase K and TE treatment at 60°C for 4 hours, and then at 98°C for 10 minutes, the DNA was first bisulfite treated (EZ DNA Methylation-Lightning Kit, Zymo Research), and then amplified and sequenced as described in ‘Targeted mutation sequencing’. Sequences were processed on Galaxy Europe (Bismark Mapper) and amplicons with incomplete conversion (C’s at non-CpG sites) were removed. After excluding 39 spots because of low epithelial content (mean methylation *β*-value of three LUMP sites <0.7), 2 spots because of low read depth (<10 target amplicon reads), and 33 spots with non-DCIS histology (benign or invasive), we analyzed a total of 68 DCIS spots across 6 tumors. Assigning each amplicon read to one of the 32 possible haplotypes (binary vectors of length 5), we visualized haplotype proportions in each spot as a measure of local epigenetic heterogeneity.

### Low-pass whole genome sequencing for spatial copy number profiling

A total of 19 spots were microdissected from DCIS-286; 9 spots corresponded to a spot with available somatic mutation calls. Whole genome libraries were prepared with NEBNext^®^ Ultra™ II DNA Library Prep Kit and sequenced on Illumina NovaSeq-6000 using paired end reads extending 150 bases and demultiplexed into pairs of FASTQ files for each sample. The FASTQ files were aligned to GRCH37 using the BWA-MEM algorithm,^48^ and the resulting BAM files were used in the CNV analysis pipeline implemented in the R package QDNAseq.^55^ Count data were obtained, smoothed, and normalized using default settings with bin annotations of size 30 kbp derived from reference genome GRCH37 as provided in the package. CNV calls were obtained using the multi-state mixture model CGHcall.^56^

### Multiplexed ion beam imaging by time of flight (MIBI-TOF)

We identified 57 fields of view (FOVs; 500*μm* x 500 *μm*) from microscope sections of 10 tumors in our cohort (range: 4 to 6 FOVs per tumor). MIBI-TOF analysis^25,57,58^ was then performed by IONPath Inc. In brief, this technology uses primary ion beam and secondary ion time-of-flight mass spectrometry to simultaneously measure protein expression and interrogate the spatial organization of tissue sections. The samples were stained with 34 metal-labeled antibodies, irradiated, and then imaged using time-of-flight mass spectroscopy. The spatial resolution of individual cells was obtained by combining the nuclear dsDNA signal with cytoplasmic and membrane markers. A deep-learning model was used to identify individual cells and score each cell for the presence of biomarkers. Cell types were determined based on the presence/absence of biomarker combinations as follows. Focusing on epithelial cells (pan cytokeratin-positive), we defined the following five epithelial cell types: luminal (BCL2-positive and/or GATA3-positive); stem-like (PAX5-positive and/or SOX10-positive); basal (CK5-positive); epithelial-to-mesenchymal (EMT; Vimentin-positive); and myoepithelial (SMA-positive). Among the 133,724 epithelial cells identified across all 57 FOVs, 78,157 (58.4%) were assigned to at least one of the 5 subtypes, and 20% (27,267) were assigned to two or more subtypes.

### Spatial registration of spots

To construct three-dimensional maps of spot locations within each tumor (**Suppl. Figure S1**), we first used the clinical pathology maps that show the spatial relationship of each paraffin block within the excised tissue. These block maps were used to locate pathologic features with respect to surgical margins and to determine the positions of each of the paraffin blocks included in the study along the long axis of the tissue/tumor (referred to as the z-axis). Once positioned along the z-axis, we oriented the thin sections from these blocks based on colored ink stains along the tissue margins. Once the slides were properly oriented, we determined the in-plane location (x- and y-coordinates) of individual spots which had been recorded during microdissection. The origin of the x- and y-coordinates were anchored at the center of each slide, and spot coordinates were recorded after accounting for microscopic magnification. Combining the in-plane x- and y-coordinates with the z-coordinate along the tumor’s long axis thus completed the process of spatial spot registration.

### Phenotypic annotation of spots

The histologic phenotype of each spot was determined in three steps. First, two board-certified breast pathologists (A.H. and J.G.) independently reviewed the H&E slides, classified each spot as ‘normal’, ‘benign’, ‘DCIS’ or ‘invasive’, and used a free text field to provide a comprehensive description of all ‘benign’ spots. Spots where the two pathologists agreed on the main category (normal, benign, DCIS, invasive) were considered complete (n=445, or 85%); the remaining spots (n=79, or 15%) were adjudicated by a third board-certified breast pathologist (D.W.). A board-certified pathologist (D.S.) used the free text annotations of all ‘benign’ spots to refine their classification as either ‘normal breast tissue,’ ‘benign breast disease without atypia’, or ‘benign breast disease with atypia’. Finally, a board-certified breast pathologist (A.H.) assigned to each DCIS spot a pathologic subtype (solid, cribriform, micropapillary) and determined whether comedo-like features were present.

### Mutation calls

Variant calling based on the tumor-specific SNV panels was performed in each spot separately, using a previously described Bayesian inference method.^59^ Briefly, for any given sequencing target, the posterior distribution of the target’s VAF *f* was calculated by combining a data likelihood and a prior distribution according to Bayes’ theorem. For the data likelihood, we used a binomial model for the variant read count *K* and the total read count *N*, accounting for a sequencing error rate *e* as follows

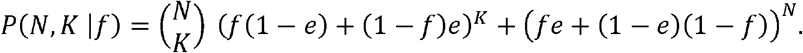

Our prior belief about the VAF was modeled as a mixture

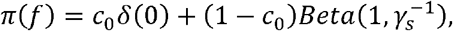

where *c*_0_ is the prior probability of the mutation being absent (as reflected by the point mass *δ*(0)), and, if present, the mutation’s VAF was assumed to have a prior distribution 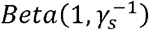 where γ*s* is the sample purity. Applying Bayes’ theorem, the posterior distribution of the VAF, or *P*(*f*|*N,K*; *e, c*_0_, *γ*_*s*_ can be calculated explicitly. Finally, the posterior probability that a mutation is absent (*q*) or present (*p*) is then given by

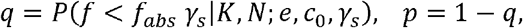

where *f*_abs_ is a pre-defined sequencing threshold. For applications where binary mutation calls were needed, we called individual SNVs absent if *q* > 95% and present if *p*> 95%. The handling of mutations with *q, p* ∈ [5%,95%] was determined *in situ*, depending on the analyses performed.

Unless otherwise specified we used the following parameter values: *e* = 0.01 which reflects the empirical error rate of the sequencing platform^60^; *f*_*abs*_ = 5% to avoid false positives mutation calls^59^; *c*_0_ = 0.5 to reflect a lack of prior knowledge about the absence vs presence of a mutation; and *γ*_s_ = 0.8 to reflect the high sample purity achieved by SURF.

### Mutational signatures

To analyze the DNA mutation patterns in our cohort, we compiled a list of targeted mutations that were present in at least one microdissected spot. Using the R package MutSignatures,^61^ we categorized mutations into 96 types based on 6 possible single base pair substitution categories (C>A, C>G, C>T, T>A, T>C and T>G) and 16 combinations of 3’ and 5’ nucleotide neighbors. We performed de novo extraction of mutational signatures using the non-negative matrix factorization method (n=1,000 bootstrap iterations, k=2 signatures) and estimated the exposure of each tumor sample to the two signatures. In separate analyses, we performed de novo extraction for k=3 and k=4 signatures; since these resulted in the same two high-quality signatures as extracted for k=2, accompanied by additional low-quality signatures, we chose k=2 for the final analysis. We then compared the two extracted signatures to the COSMIC database (https://cancer.sanger.ac.uk/cosmic) using the cosine distance, and further assessed whether matching signatures were breast cancer related^62^ or possible sequencing artefacts.

### Driver mutation annotation

Single nucleotide variants were annotated using the SIFT annotation tool (https://sift.bii.a-star.edu.sg/), which predicts mutation effect and functional impact on the protein. Briefly, SNVs were organized into a VCF file format, specifying chromosome, genomic position, and reference and alternate alleles according to the GCRh37. The VCF file was input into the SIFT Java executable tool, which output annotated SNVs, labeling Ensembl transcript and gene IDs, gene name, coding region (CDS, UTR_3, UTR_5), variant type (noncoding, nonsynonymous, stop-gain, substitution, synonymous), and functional prediction (deleterious, tolerated). SNVs with no gene label were labeled as intergenic, and SNVs with a gene label but not in a coding or UTR region were labeled as intronic. Protein coding changes in genes that have been functionally associated with breast cancer in either the TCGA (https://www.cancer.gov/tcga) or COSMIC (https://cancer.sanger.ac.uk/cosmic) databases were considered putative driver mutations (**Suppl. Table S2**). All others were categorized as passenger mutations.

### Final study cohort

After eliminating two tumors due to PCR issues, the remaining 19 tumors comprised a total of 524 individual spots and 1,108 targeted loci. Among the 31,265 sequencing targets (each target is a spot-SNV pair), there were 7,130 (22.8%) low-quality targets (LQTs) where either no sequencing results were obtained or the total absolute read count was less than 20. After removing 6 spots with undefined histology, 247 mutations that constituted LQTs in more than 40% of assayed spots, 32 germline mutations (which were present in the matched normal with a probability ≥99%), and an additional 45 spots that contained more than 40% of LQT, there were fewer than 5% of LQTs left among the 22,612 targets. At this stage, we excluded one more tumor (DCIS-221) because of low information content: 11 of the 14 detected mutations were germline mutations, and the remaining 3 mutations were detected in only one spot each. An overview of the LQT removal process among the 18 tumors included in the final study cohort is found in **Suppl. Figure S2**.

### Uncertainty quantification

For tumor statistics based on binary mutation calls, we leveraged the Bayesian framework to propagate posterior uncertainty through Monte Carlo sampling. More precisely, for a tumor with *N* spots and *M* mutations, we sampled *T* independent and identically distributed binary spot-mutation arrays *S* = (*s*_*ij*_) ∈ ℝ^*N* × *M*^, where *s*_*ij*_ ∼ *Bernoulli*(*p*_*ij*_) and *p*_*ij*_ is the posterior probability of mutation *j* being present in spot *i*. For LQTs, because there was no data available, we used the prior probability instead. The statistic of interest was then computed for each of the *T* realizations of *S*, and the posterior predicted mean and 95% prediction interval were recorded. Unless otherwise specified, the default was *T* = 1,000.

### Spatial-genetic correlation

A tumor-level spatial-genetic correlation measure was introduced to assess the degree of spatial intratumor heterogeneity. Uncertainty was quantified as described above, and we focus here on the derivation of the statistic for a single realization of the binary array *S* = (*s*_*ij*_) ∈ ℝ^*N* × *M*^, for a tumor with *N* spots and *M* mutations. First, we defined the spatial distance *d*_*s*_ (*i, j*) between two spots *i* and *j* as their Euclidian (*L*^2^) distance in ℝ^3^. Next, we introduced the notion of spot *i*’s genotype as the vector *g*_*i*_ = (*s*_*i1*_, *s*_*i2*_, *s*_*iM*_) ∈ ℝ^*M*^ and defined the genetic distance *d*_*g*_ (*i, j*) between two spots *i* and *j* as the Manhattan (*L*^1^) distance between *g*_i_ and *g*_j_. Finally, we calculated the spatial and genetic distances between all *N* (*N* − 1) spot pairs and computed their correlation (Pearson’s R).

### Expansion index

The expansion index (EI) was introduced to distinguish, at the tumor level, between spatially discontinuous ‘skip’ lesions and continuous ‘patch’ lesions (**Suppl. Figure S6**). Again, uncertainty quantification was performed as described above, and we focus here on the derivation of the statistic for a single realization of the binary array *S* = (*s*_*ij*_) ∈ ℝ^*N* × *M*^, for a tumor with spots *N* and *M* mutations. The definition of the EI is based on a bivariate characterization 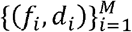 of the tumor’s mutations, where *f*_i_ is the fraction of DCIS spots in which mutation *i* is present, and *d*_i_ is the normalized diameter of mutation *i*, defined as the maximum Euclidian distance between any two DCIS spots containing the mutation, divided by the maximum Euclidian distance between any two DCIS spots in the tumor. As illustrated in **Suppl. Figure S6**, the *EI* is then obtained by integrating the piecewise constant curve over the *M* bivariate points

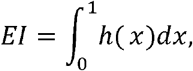

where

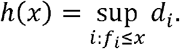

By definition, *EI* ∈ [0,1]. If mutation diameter grows approximately linearly with the fraction of occupied spots, then *EI* ∼ 0.5, indicative of a continuous patch lesion. If there are mutations with a large diameter at a low fraction of occupied spots, *EI* » 1, indicative of a disperse skip lesion.

### Mutation energy

This statistic was introduced to quantify the mutational diversity of the tumor. Again, uncertainty quantification was performed as described above, and we focus here on the derivation of the statistic for a single realization of the binary array *S* = (*s*_*ij*_) ∈ ℝ^*N* × *M*^, for a tumor with *N* spots and *M* mutations, each of which was detected in ≥1 spot(s). First, we applied hierarchical column clustering (using the Manhattan distance) to obtain the spot genotype-clustered array 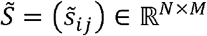. Next, in analogy with the Ising model from statistical mechanics,^63^ we defined the mutation energy *I*_*k*_ of mutation *k* as

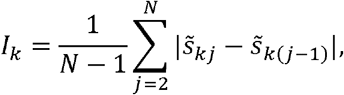

where the normalizing factor accounts for the *N* − 1 possible flips and ensures that *I*_*k*_ ∈ [0,1] irrespective of the number of spots in the tumor. Intuitively, *I*_*k*_ measures, for each mutation, the number of “flips” from “absent” to “present” along the rows of the spot-mutation array (e.g., **Suppl. Figure S3**). If there is only a single spot genotype in the tumor, then *I*_*k*_ *= 0* for all *k*; and *I*_*k*_ increases as both the number of different spot genotypes and their degree of dissimilarity increase. To quantify the mutation energy at the tumor rather than individual mutation level, we used the median and interquartile range (IQR) across all detected mutations in the tumor.

### Dimension reduction of genotype space

First, we assigned each spot *i* (*i* = 1, …, *N*) a vector of posterior mutation probabilities *p* = (*p*_*ij*_) ∈ ℝ^*M*^, where *p*_*ij*_ is the posterior probability of mutation *j* being present in spot *i*, and *M* is the number of detected mutations in the tumor. For low-quality targets and targets with read count <20, *p*_*ij*_ was set to the prior probability of the mutation being detected. Next, we applied t-distributed stochastic neighbor embedding to reduce the genotype space to two dimensions (package *Rtsne*, v0.16, with perplexity= (*N* − 1) /3 and default settings otherwise).

### Multiclonal invasion

For synchronous DCIS tumors, multiclonal invasion is defined as the co-occurrence of 2 or more subclones that are present in both the *in situ* and invasive compartments of the tumor.^11^ Because subclone deconvolution is not practical for SNV panels of limited size, we derived a necessary and sufficient condition for multiclonality in terms of individual mutations as follows. We identified mutations that were restricted (present in <90% of eligible spots) in both the DCIS and invasive portions of the tumor. If a mutation satisfies this pattern, this implies the existence of at least two distinct subclones (one with and one without the mutation) both of which are present in the DCIS and invasive tumor portions, thus satisfying the definition of multiclonal invasion. For each tumor, we counted the number of such unique mutation patterns.

### Mathematical models of DCIS growth

Here, we provide a summary only; details about model formulation and model fitting are found in the **Technical Appendix**. To model DCIS growth, we combined a generative stochastic model of the binary ductal tree structure with a stochastic model of the cellular DCIS growth dynamics. The ductal tree model was based on the experimentally delineated dynamics of branching ductal morphogenesis, that is ductal elongation followed by either branching into two daughter ducts, or branch termination, with equal probability.^15^

Tumor growth along the ductal tree architecture was initiated by random seeding of the first DCIS cell. Growth from this first cell to the macroscopic tumor was modeled as a two-stage process, consisting of an initial exponential expansion subject to the mutation bursts of punctuated evolution (with mutation rate per cell division), followed by an expansive growth along the branching tree structure. To describe the expansive growth phase, we considered three competing models as follows.

*Model 1*, or Comet model, recapitulates the cellular dynamics of pubertal branching morphogenesis of the mammary gland.^15,17^ In this model, the DCIS end buds are nucleated by a pool of *N* long-lived cancer cells, half of which undergo intermittent asymmetric division followed by *n*_TA_ generations of transit-amplification, and half of which remain quiescent. As the end buds of the growing tumor thus move along the pre-existing ducts, the transit-amplifying progenies of the dividing end bud cells contribute to the growing tumor. Upon reaching a ductal branching point, the long-lived end bud cells are randomly divided between the two daughter branches, and after a round of duplication, the two newly created end buds begin to grow along the respective daughter ducts.

*Model 2* is a variation of Model 1, whereby all cells in the DCIS end bud are assumed to undergo intermittent asymmetric division. This variation of the Comet model was introduced to assess its sensitivity to the separation of proliferating and quiescent end bud cells.

*Model 3* is a canonical cancer evolution model characterized by uncontrolled proliferation and competition among DCIS cells. To account for spatial crowding and resource constraints behind the actively growing tips of the tumor, we formulated a boundary growth model where only the *N* cells immediately behind the growing tips contribute to the net growth of the elongating DCIS duct. The same branching dynamics as in Models 1 and 2 were applied.

### Model fitting and model selection

We used a rejection sampling-based version of approximate Bayesian computation to fit the models to the experimental data, estimate the posterior parameter distributions (*N*, μ, *n*_TA_ ) and identify the best fitting model.^64,65^ For a given model, we sampled a set of parameters from the prior distributions (see **Technical Appendix**), simulated a ductal tree and DCIS tumor as described above, and compared the simulated tumor against the experimental tumors in our study cohort using a distance function. By keeping only parameter sets resulting in simulated tumors that were sufficiently similar to the experimental data—that is, the distance between simulation and experiment was below a specified threshold—we thus approximated the posterior parameter distributions. Finally, we used a joint model-parameter space approach^66^ to compute the posterior marginal model probabilities and calculate the Bayes’ factors for model selection.

### Statistical analyses

All statistical analyses were performed using R version 4.2.0 (R Foundation for Statistical Computing, Vienna, Austria). All statistical tests were two-sided. Data visualizations were made with R, using the packages *ggplot2* (v3.3.6), *ggbeeswarm* (v0.6.0), and *circlize*^*67*^ (v0.4.16).

## DATA AVAILABILITY

Upon publication of the manuscript, the whole exome and targeted sequencing data will be deposited in the Sequence Read Archive database (https://www.ncbi.nlm.nih.gov/sra) under a unique accession code. All other data supporting the findings of this study are available within the paper and its supplementary files or available from the corresponding authors upon reasonable request.

## CODE AVAILABILITY

The code archive has been submitted alongside the manuscript. Upon publication of the manuscript, the code used to produce the results in this manuscript will be made available at https://github.com/mdryser/D5_DCIS (MIT License).

## ACKNOWLEDGMENTS

We gratefully recognize our funders who provided support for this work: National Institutes of Health (grant K99-CA207872 to M.D.R.; grant R00-CA207872 to M.D.R.; grant U2C-CA233254 to E.S.H. and C.C.M.; grant R01-CA185138 to E.S.H., grant U54-CA217376 to C.C.M. and D.S., grant P01-CA91955 to C.C.M.; grant R01-CA140657 to C.C.M.; and grant U01-CA214183 to J.R.M.), National Science Foundation (grant DMS-1614838 to M.D.R.), Department of Defense (grant BC132057 to E.S.H), Breast Cancer Research Foundation (grant BCRF-19-074 to E.S.H), CDMRP Breast Cancer Research Program (grant BC132057 to C.C.M.), and Arizona Biomedical Research Commission (grant ADHS18-198847 to C.C.M.).

## AUTHORSHIP CONTRIBUTIONS

M.D.R., D.S., and E.S.H. designed and managed the study; M.D.R., A.H., L.M.K., J.R.M., and E.S.H. identified the tumor samples; L.M.K. managed IRBs; A.H., L.J.G., D.S., and E.S.H. contributed clinical expertise; A.H., J.G., D.L.W., and D.S. performed pathology reviews; M.D.R., I.C.S., E.W., and D.S. performed mathematical modeling; D.M., C.C.M., J.R.M., and E.S.H. performed WES; M.D.R. and D.S. designed targeted sequencing panels; D.S. performed tissue microdissection and targeted sequencing; K.M. and J.R.M. annotated the mutations, M.D.R., M.G., I.C.S., and D.S. analyzed the data; M.D.R. and M.G. visualized the data; M.D.R., M.G., J.R.M., D.S., and E.S.H. wrote the manuscript; all authors provided feedback throughout the study and reviewed the final manuscript.

## OVERVIEW OF SUPPLEMENTARY MATERIALS

**Technical Appendix:** Mathematical modeling and statistical inference

**Supplementary Table S1**: Study cohort overview

**Supplementary Table S2**: Putative driver mutations

**Supplementary Table S3:** Marginal model probabilities and Bayes’ factors

**Supplementary Table S4:** Invasion analysis among 8 synchronous DCIS

**Supplementary Table S5:** Overview of model parameters

**Supplementary Figure S1:** Spatial spot maps

**Supplementary Figure S2:** Removal of low-quality targets in final study cohort

**Supplementary Figure S3:** Tumor summaries

**Supplementary Figure S4:** Mutational signatures

**Supplementary Figure S5:** Spatially clustered tumor summaries

**Supplementary Figure S6:** Expansion index

**Supplementary Figure S7:** VAF spectra by mutation subgroups

**Supplementary Figure S8:** Mutation diameter by driver status

**Supplementary Figure S9:** Spatial copy number analysis

**Supplementary Figure S10:** Model fits

**Supplementary Figure S11:** Targeted methylation sequencing

**Supplementary Figure S12**: Phenotype-genotype analysis I

**Supplementary Figure S13:** Phenotype-genotype analysis II

**Supplementary Figure S14:** Model framework

**Supplementary Figure S15:** Stochastic tree model

**Supplementary Figure S16:** DCIS ductal elongation and branching dynamics

